# Differentiation of human adult-derived stem cells towards a neural lineage involves a de-differentiation event prior to re-differentiation to neural phenotypes

**DOI:** 10.1101/2021.02.04.429708

**Authors:** Carlos Bueno, Marta Martínez-Morga, David García-Bernal, José M Moraleda, Salvador Martínez

## Abstract

Although it has been reported that mesenchymal stem cells isolated from adult tissues can be induced to overcome their mesenchymal fate and transdifferentiate into neural cells, the findings and their interpretation have been challenged. The main argument against this process is that the cells rapidly adopt neuron-like morphologies through retraction of the cytoplasm rather than active neurite extension.

In this study, we examined the sequence of biological events during neural differentiation of human periodontal ligament-derived stem cells (hPDLSCs), human bone marrow-derived stem cells (hBMSCs) and human dental pulp-derived stem cells (hDPSCs) by time-lapse microscopy.

We have demonstrated that hPDLSCs, hBMSCs and hDPSCs can directly differentiate into neuron-like cells without passing through a mitotic stage and that they shrink dramatically and change their morphology to that of neuron-like cells through active neurite extension. Furthermore, we observed micronuclei movement and transient cell nuclei lobulation concurrent to *in vitro* neurogenesis from hBMSCs and hDPSCs.

Our results demonstrate that the differentiation of hPDLSCs, hBMSCs and hDPSCs towards a neural lineage occurs through a de-differentiation step followed by re-differentiation to neural phenotypes, and therefore we definitively confirm that the rapid acquisition of the neural phenotype is via a differentiation trait.

## Introduction

In the classical view of cell development, embryonic stem cells differentiate into progressively more precursor cells with an increasingly restricted lineage potential, until the final mature, specialised cell types are generated and functionally integrated into their respective tissues. The differentiated state of a cell was believed to be terminal and irreversible^1,2^. While asymmetric cell division is considered to be the mechanism by which the asymmetric inheritance of cellular components during mitosis defines the distinct fate of each daughter cell^3^.

However, there is increasing evidence that the rules of irreversibility and germ-layer restriction can be broken^4^. It has long been accepted that cells can change their identity, both *in vitro* and *in vivo*, a phenomenon known as cellular plasticity^5,6^. Cellular conversion falls into two major categories: de-differentiation and transdifferentiation^4-8^. De-differentiation is the conversion from one differentiated cell stage to a less differentiated stage within the same lineage. Transdifferentiation is the conversion of one differentiated cell type into another, either within or across germ layers. It has been suggested that transdifferentiation may involve a de-differentiation step before cells re-differentiate to a new mature phenotype, or it may occur directly in a process that bypasses such immature phenotypes^9^.

Literature published over the past 20 years have shown that mesenchymal stem cells (MSCs) isolated from adult tissues can be induced to overcome their mesenchymal fate and transdifferentiate into neural cells^10-17^. Although the transdifferentiation of human MSCs into neurons has aroused considerable interest, as it would have immense clinical potential in cell replacement therapy and regenerative medicine^18,19^, the findings and their interpretation have been challenged^20,21^. The main argument against this process is that the cells rapidly adopt neuron-like morphologies through retraction of the cytoplasm rather than by active neurite extensión^22,23^.

In a recent study^24^, we showed that human periodontal ligament-derived stem cells (hPDLSCs) can directly differentiate into neuron-like cells without passing through any mitotic stages. When hPDLSCs were exposed to a neural induction medium, we found that they rapidly underwent a dramatic change in shape and size; they initially adopted highly irregular forms before gradually contracting into round cells (neurogenesis). These round cells then slowly adopted a complex morphology and finally gave rise to a variety of neuron-like morphologies (neuronal polarisation). Furthermore, we also reported nuclear remodelling concurrent to *in vitro* neurogenesis from hPDLSCs.

The present study is designed to confirm that human stem cells isolated from adult tissues in the early stages of exposure to a neural induction medium adopt a neuron-like morphology because of a differentiation trait rather than an artefact. We performed time-lapse phase-contrast microscopy to record the changes in the cell morphology of hPDLSCs, human bone marrow-derived stem cells (hBMSCs) and human dental pulp-derived stem cells (hDPSCs).

We found that hPDLSCs, hBMSCs and hDPSCs can directly differentiate into neuron-like cells without passing through a mitotic stage. All three types of stem cell shrunk dramatically and changed their morphology to that of neuron-like cells with active neurite extension. Additionally, we may have evidenced micronuclei movement and transient cell nuclei lobulation in parallel with *in vitro* neurogenesis from hBMSCs and hDPSCs. Our results clearly indicate that the differentiation of hPDLSCs, hBMSCs and hDPSCs towards a neural lineage occurs through a de-differentiation step before the cells re-differentiate to neural phenotypes. These findings definitively establish that the rapid acquisition of a neuron-like morphology during neural differentiation is due to a differentiation trait as opposed to an artefact.

## Results

### Time-lapse microscopy of hPDLSCs cultured in neural induction media

To conclusively affirm that hPDLSCs adopt a neuron-like morphology in the early stages of exposure to a neural induction medium is due to active neurite extension rather than retraction of the cytoplasm, we performed time-lapse phase-contrast microscopy and immunocytochemical analysis within the first 24 h of neural differentiation.

Time-lapse imaging revealed that, after neural induction, the hPDLSCs underwent a rapid change in shape and size, first adopting highly irregular forms and then gradually contracting into round cells. Subsequently, we observed the growth of new neurites from the cell body of round cells (Fig. 1, black arrows). The round cells were also noted to gradually adopt a complex morphology, acquiring dendrite-like (Fig. 2a, white arrows) and axon-like identities (Fig. 2a, red arrows), giving rise to a variety of neuron-like morphologies (Figs. 1 and 2).

**Figure 1.**
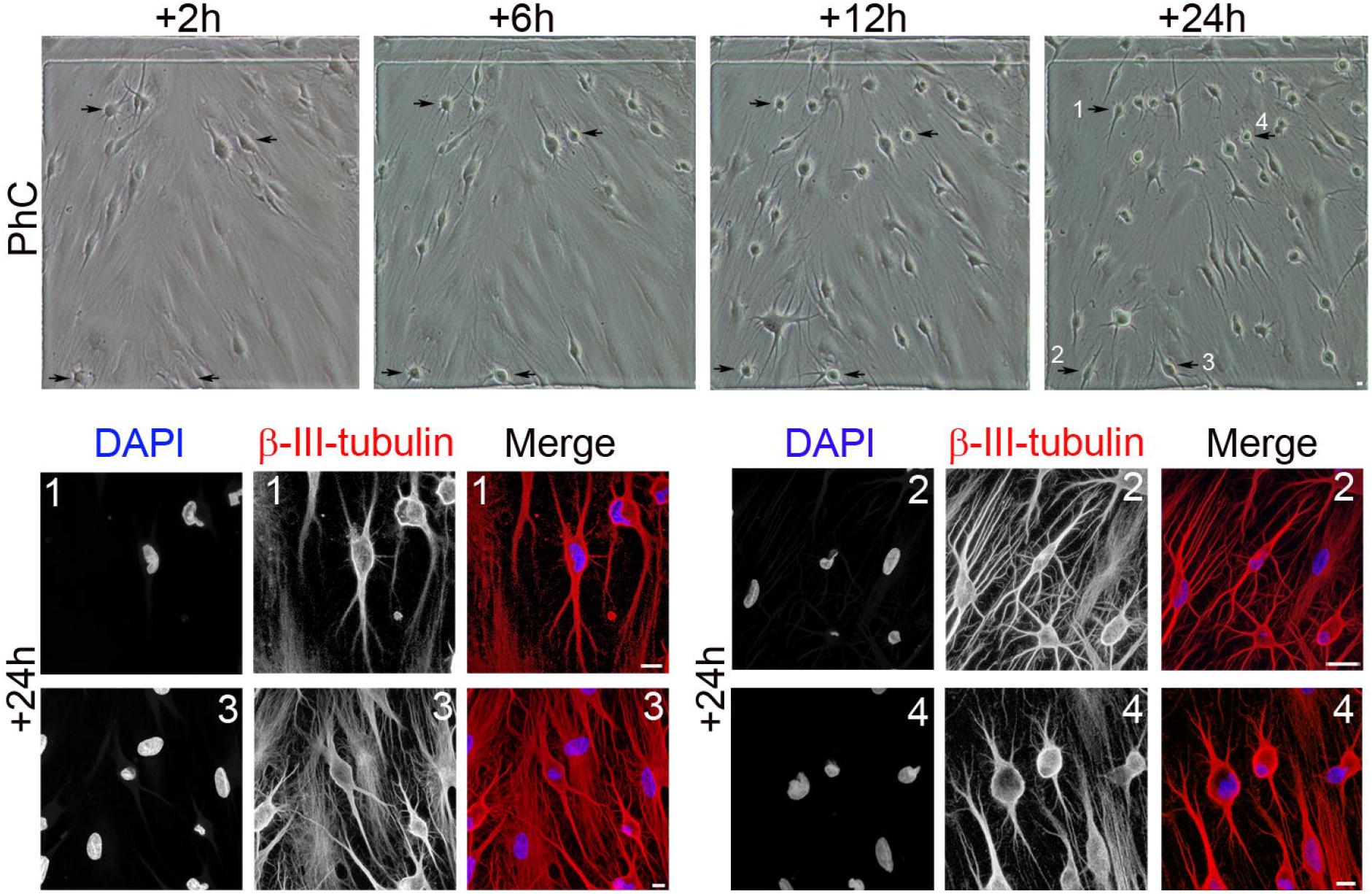
Morphological changes in hPDLSC cultures during neural induction. Time-lapse imaging and immunocytochemical analysis revealed that, after neural induction, hPDLSCs underwent a rapid change in shape and size, first adopting highly irregular forms and then gradually contracting into round cells. Subsequently, there was growth of new neurites from the cell body of round cells (black arrows) (numbers correspond to the areas shown in higher magnifications). Scale bar: 10 μm. PhC: Phase-contrast photomicrographs.

**Figure 2.**
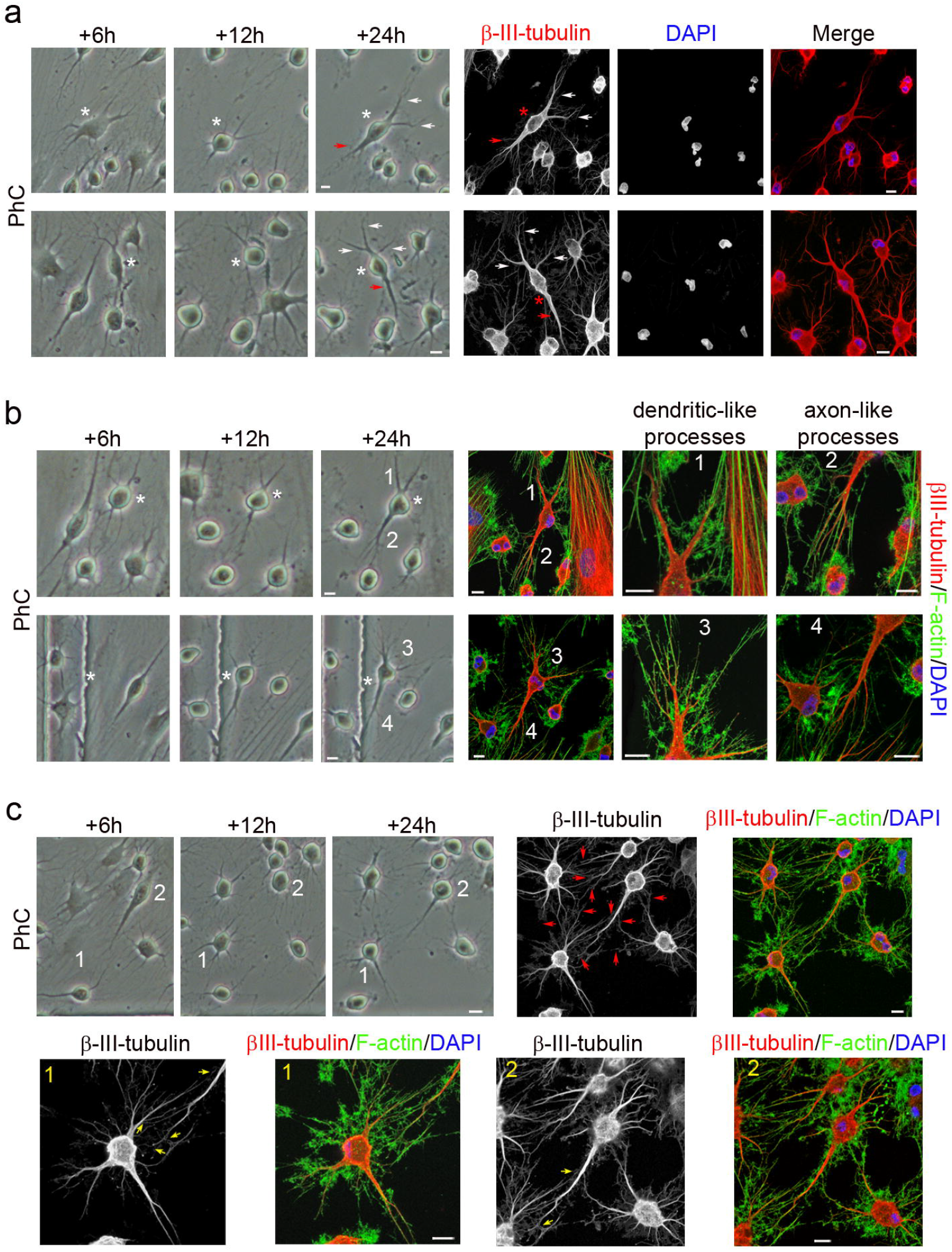
Neuronal polarisation of hPDLSC-derived neuron-like cells. (**a**) Time-lapse imaging and immunocytochemical analysis revealed that round cells gradually adopted a complex morphology, acquiring dendrite-like (white arrows) and axon-like identities (red arrows). (**b**) Cytoskeletal protein β-III tubulin and F-actin staining showed that hPDLSC-derived neuron-like cells developed well-differentiated dendrite-like and axon-like domains (numbers correspond to the areas shown in higher magnifications). (**c**) Time-lapse imaging and immunocytochemical analysis also revealed that the hPDLSC-derived neuron-like cells connected together through different types of synapse-like interactions (red arrows), including axodendritic-like synapses (yellow arrows) (numbers correspond to the areas shown in higher magnifications). Scale bar: 10 μm. PhC: Phase-contrast photomicrographs.

In addition, cytoskeletal protein β-III tubulin and F-actin staining showed that hPDLSC-derived neuron-like cells developed well-differentiated dendrite-like and axon-like domains (Fig. 2b). Time-lapse imaging revealed that the hPDLSC-derived neuron-like cells connected to one another through different types of synapse-like interactions (Fig. 2c, red arrows), including axodendritic-like synapses (Fig. 2c, yellow arrows).

hPDLSCs did not differentiate at the same time, so the cell culture simultaneously contained hPDLSCs at different stages of neurogenesis and neuronal polarisation. The results demonstrate that hPDLSCs can directly differentiate into neuron-like cells without passing through a mitotic stage, thus definitively confirming that the rapid acquisition of a neuron-like morphology during neural differentiation is by means of a differentiation trait rather than merely an artefact.

### Time-lapse microscopy of hBMSCs and hDPSCs cultured in neural induction media

Isolation, characterisation and neural differentiation of multipotent stem cells from human bone marrow^10,18,19,25,26^ and dental pulp^27-30^ have been described previously. Under proliferation conditions, hBMSCs and hDPSCs displayed a fibroblast-like morphology with actin microfilaments and β-III tubulin microtubules oriented parallel to the cell’s longitudinal axis (Fig. S1). During interphase, undifferentiated hBMSCs and hDPSCs had an ellipsoidal nucleus, often located in the centre of the cell (Fig. S1). These results are consistent with previous studies that found hBMSCs and dental-derived stem cells exhibited spontaneous expression of neural marker β-III tubulin even without neural induction^31^.

To determine whether the same rapid morphological changes could also be induced in other human adult-derived stem cells, we tested hBMSCs and hDPSCs under the same neural induction conditions. As with hPDLSCs, time-lapse imaging revealed that after neural induction hBMSCs and hDPSCs underwent a dramatic change in shape and size, first adopting highly irregular forms, before gradually contracting into round cells. Subsequently, we noted the growth of new neurites from the cell body of hBMSC-derived (Fig. 3a, black arrows) and hDPSC-derived round cells (Fig. 3b, black arrows).

**Figure 3.**
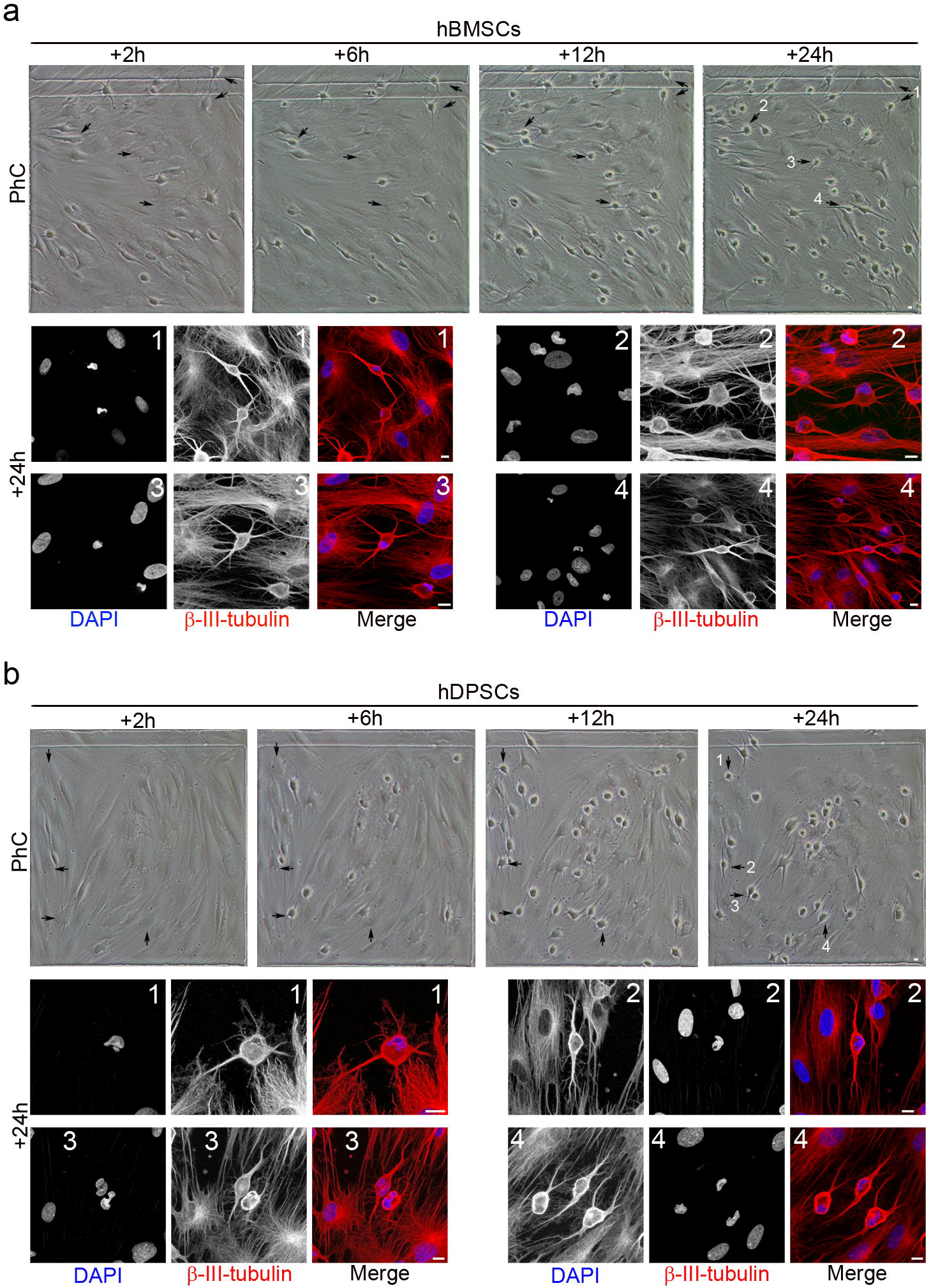
Morphological changes in hBMSC and hDPSC cultures during neural induction. Time-lapse imaging and immunocytochemical analysis revealed that, after neural induction, hBMSCs (**a**) and hDPSCs (**b**) underwent a rapid change in shape and size, first adopting highly irregular forms and then gradually contracting into round cells. Subsequently, there was growth of new neurites from the cell body of round cells (black arrows) (numbers correspond to the areas shown in higher magnifications). Scale bar: 10 μm. PhC: Phase-contrast photomicrographs.

We also observed that hBMSC-derived (Fig. 4a) and hDPSC-derived neuron-like cells (Fig. 4b) gradually adopted a complex morphology, acquiring dendrite-like (Fig. 4, white arrows) and axon-like identities (Fig. 4, red arrows), and giving rise to a variety of neuron-like morphologies (Figs. 3-5). Cytoskeletal protein β-III tubulin and F-actin staining showed that hBMSC-derived (Fig. 5A) and hDPSC-derived neuron-like cells (Fig. 5b) developed well-differentiated dendrite-like and axon-like domains. Morphological analysis also revealed that hBMSC-derived neuron-like cells were interconnected by different types of synapse-like interactions, including dendrodendritic-like synapses (Fig. 5c, red arrows) and axoaxonic-like synapses (Fig. 5c, yellow arrows).

**Figure 4.**
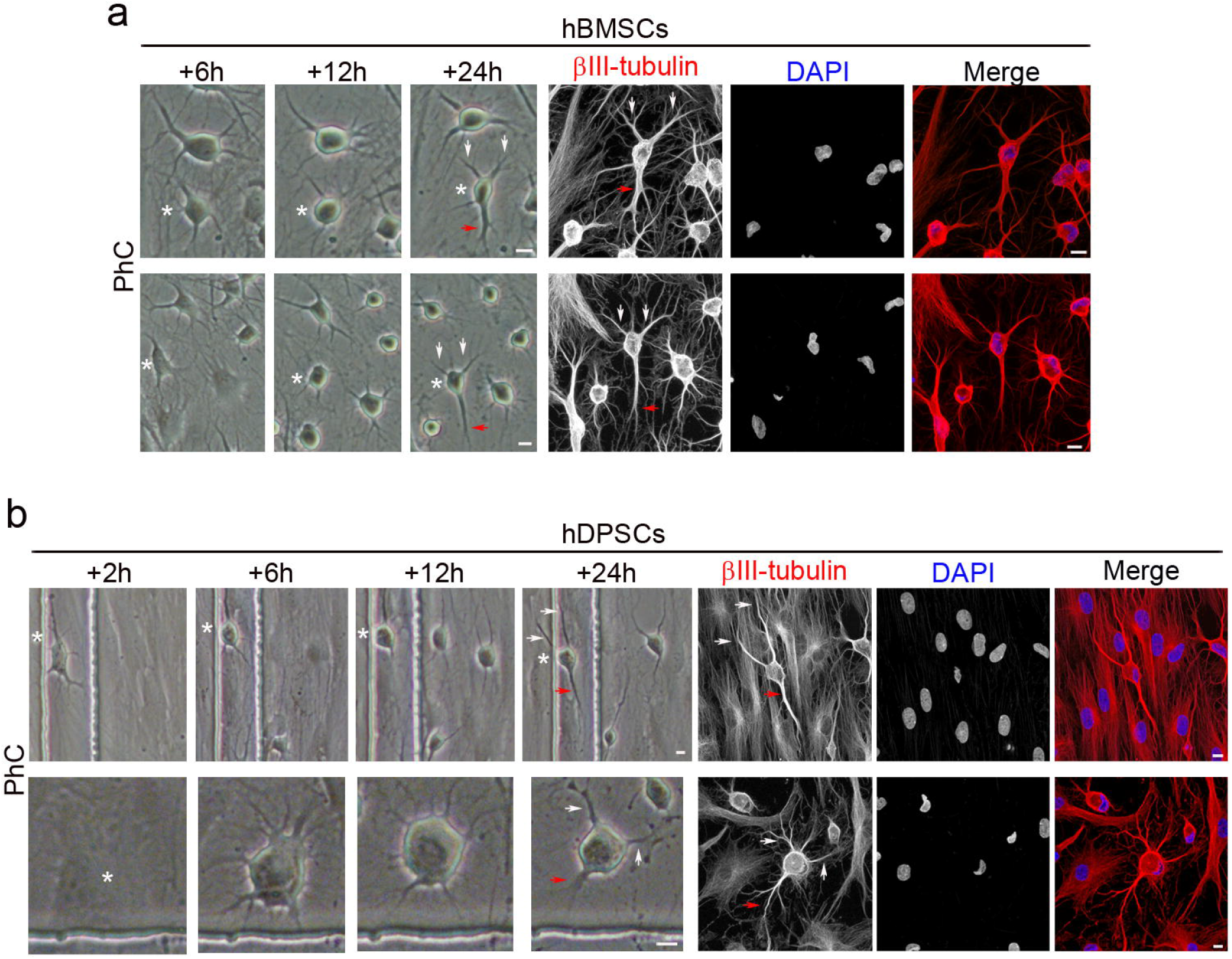
Early stages of neuronal polarisation of hBMSC-derived and hDPSC-derived neuron-like cells. Time-lapse imaging and immunocytochemical analysis revealed that hBMSC-derived (**a**) and hDPSC-derived neuron-like cells (**b**) gradually adopted a complex morphology, acquiring dendrite-like (white arrows) and axon-like identities (red arrows). Scale bar: 10 μm. PhC: Phase-contrast photomicrographs.

**Figure 5.**
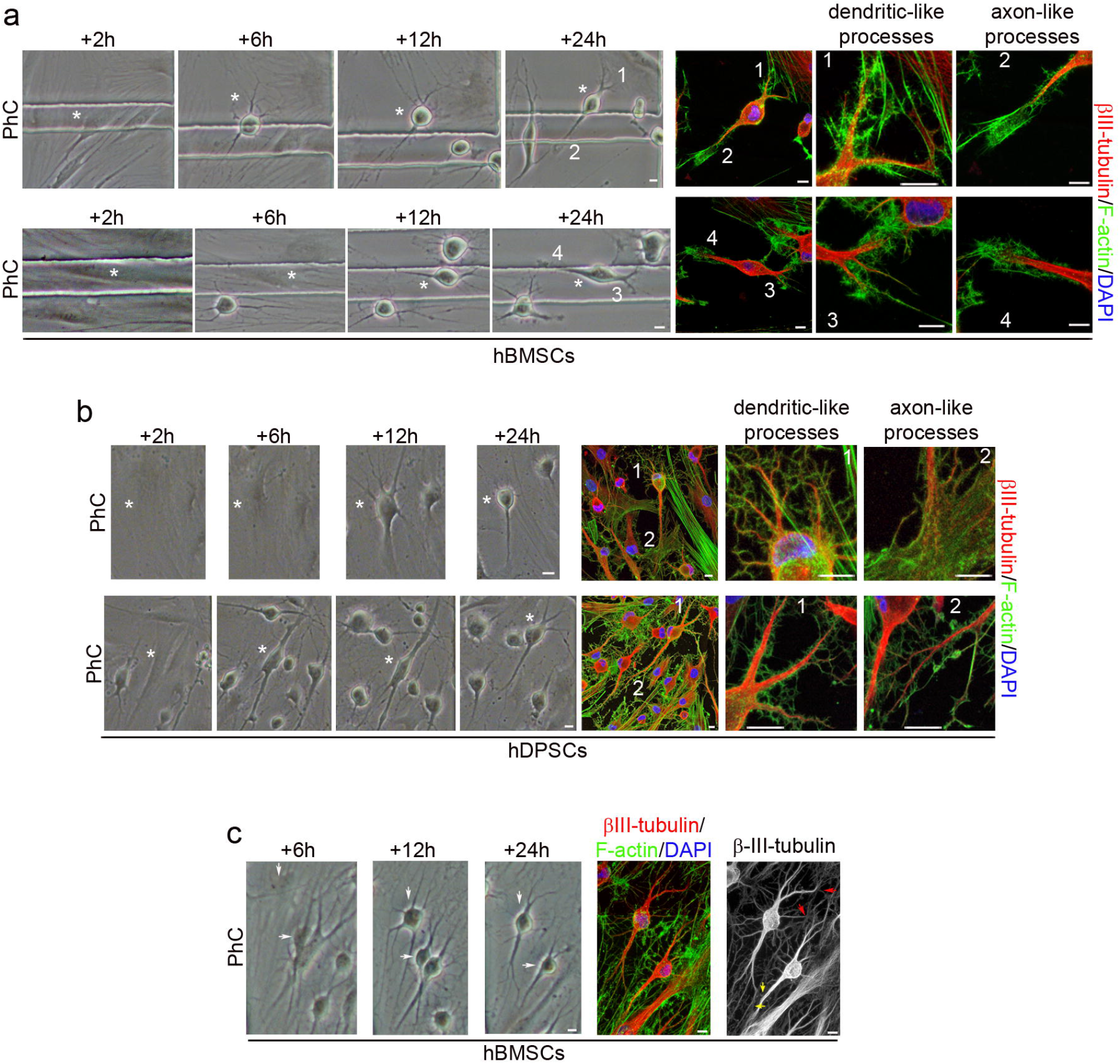
Final stages of neuronal polarisation of hBMSC-derived and hDPSC-derived neuron-like cells. Cytoskeletal protein β-III tubulin and F-actin staining showed that hBMSC-derived (**a**) and hDPSC-derived neuron-like cells (**b**) developed well-differentiated dendrite-like and axon-like domains (numbers correspond to the areas shown in higher magnifications). (**c**) Time-lapse imaging and immunocytochemical analysis also revealed that the hBMSC-derived neuron-like cells connected together through different types of synapse-like interactions, including dendrodendritic-like synapses (red arrows) and axoaxonic-like synapses (yellow arrows). Scale bar: 10 μm. PhC: Phase-contrast photomicrographs.

The results indicate that there was no cell proliferation during neurogenesis from hBMSCs and hDPSCs. The undifferentiated spindle-shaped cells were reset and started their neuronal development as round spheres. We observed that the hBMSCs and hDPSCs did not differentiate at the same time and therefore the cell culture simultaneously contained hBMSCs and hDPSCs at different stages of neurogenesis and neuronal polarisation.

These findings indicate that hPDLSCs, hBMSCs and hDPSCs have similar morphological neural development sequences *in vitro*. The differentiation of hPDLSCs, hBMSCs and hDPSCs towards a neural lineage occurs through a de-differentiation step before the cells re-differentiate to neural phenotypes.

### Nuclear remodelling

In a recent study^24^, we showed that nuclear remodelling occurred during *in vitro* neurogenesis from hPDLSCs. We discovered that many hPDLSCs exhibit unusual nuclear structures and extranuclear bodies in the cellular cytoplasm when hPDLSCs approach a spherical morphology (neurogenesis). In addition, no unusual nuclear structures were observed as hPDLSC-derived neuron-like cells gradually acquired a more mature neuron-like morphology (neuronal polarisation).

While we acknowledge that the definitive nuclear remodelling sequence, when hPDLSCs become near-spherical, can only be determined by detecting fluorescent labelled cell nuclei in time-lapse microscopy, our accumulated data suggest how these steps may occur^24^. Chromatin-containing bodies arise from the main nuclei and start to move towards specific positions within the cell, temporarily forming lobed nuclei. Subsequently, these lobed nuclei connect together to form nucleoplasmic bridges and finally the internuclear bridges facilitate the merger into a single nucleus with an eccentric position within the cell.

To determine whether nuclear remodelling also occurs when hBMSCs and hDPSCs take on a near-spherical shape, we differentiated hPDLSCs, hBMSCs and hDPSCs in the same neural induction medium (Figs. 6-8).

**Figure 6.**
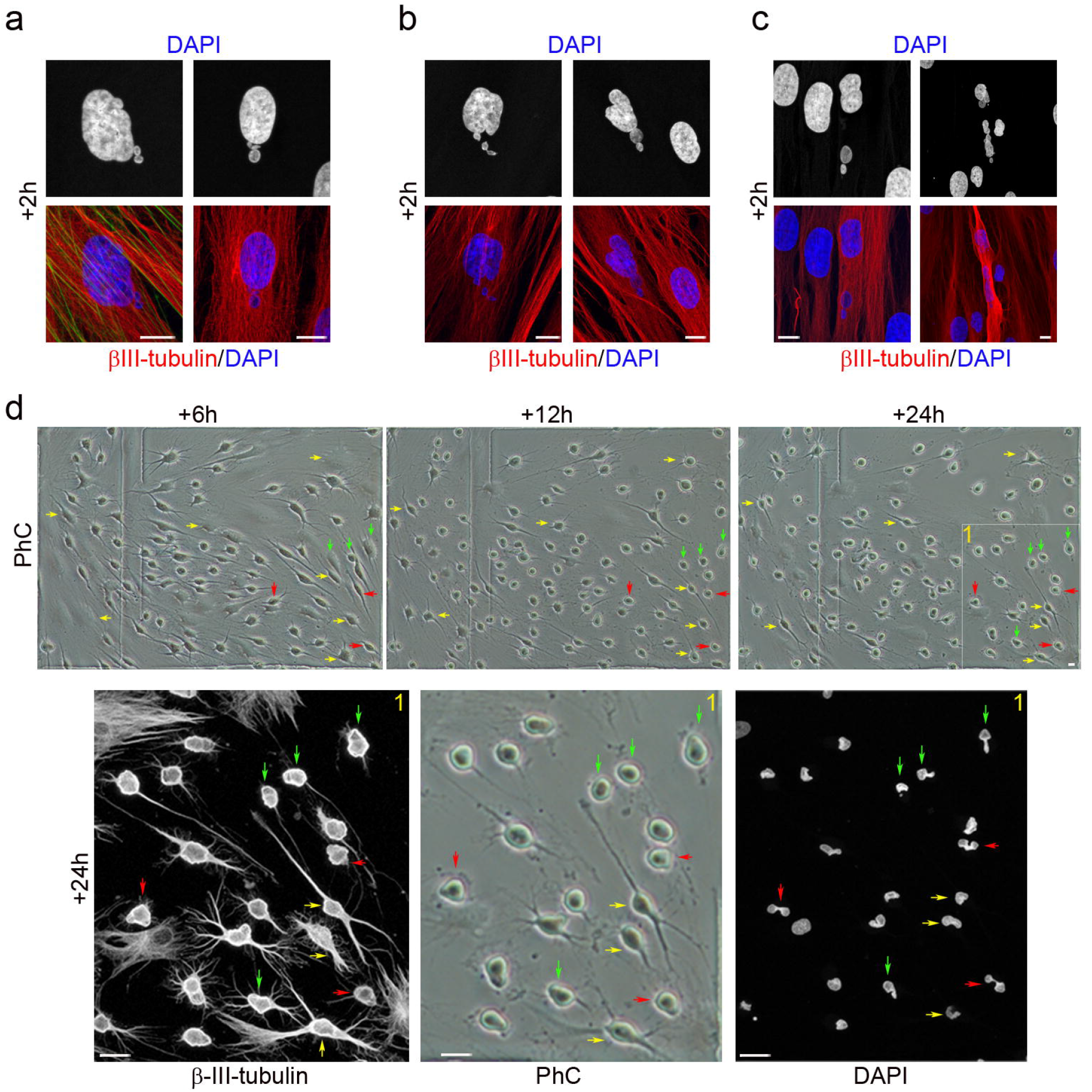
Nuclear shape remodelling occurs during neurogenesis from hPDLSCs. DNA-containing structures arose from the main nuclei (**a**) and started to move (**b**) towards specific positions within the cell (**c**), and temporarily form lobed nuclei. These lobed nuclei then connected to one another through small DNA-containing structures to form nucleoplasmic bridges (**d**, red arrows). Finally, the lobed nuclei connected by internuclear bridges (**d**, green arrows) joined to form single nuclei with an eccentric position within hPDLSC-derived neuron-like cells (**d**, yellow arrows). Scale bar: 10 μm. PhC: Phase-contrast photomicrographs.

Within 2 h of exposure to the neural induction medium we observed DNA-containing structures arise from the main nuclei of hPDLSCs (Fig. 6a), hBMSCs (Fig. 7a) and hDPSCs (Fig. 8a). We also observed DNA-containing structures moving away from the main nuclei of and towards specific positions within hPDLSCs (Figs. 6b and 6c, respectively), hBMSCs (Figs. 7b and 7c) and hDPSCs (Figs. 8b and 8c), where they temporarily formed lobed nuclei.

**Figure 7.**
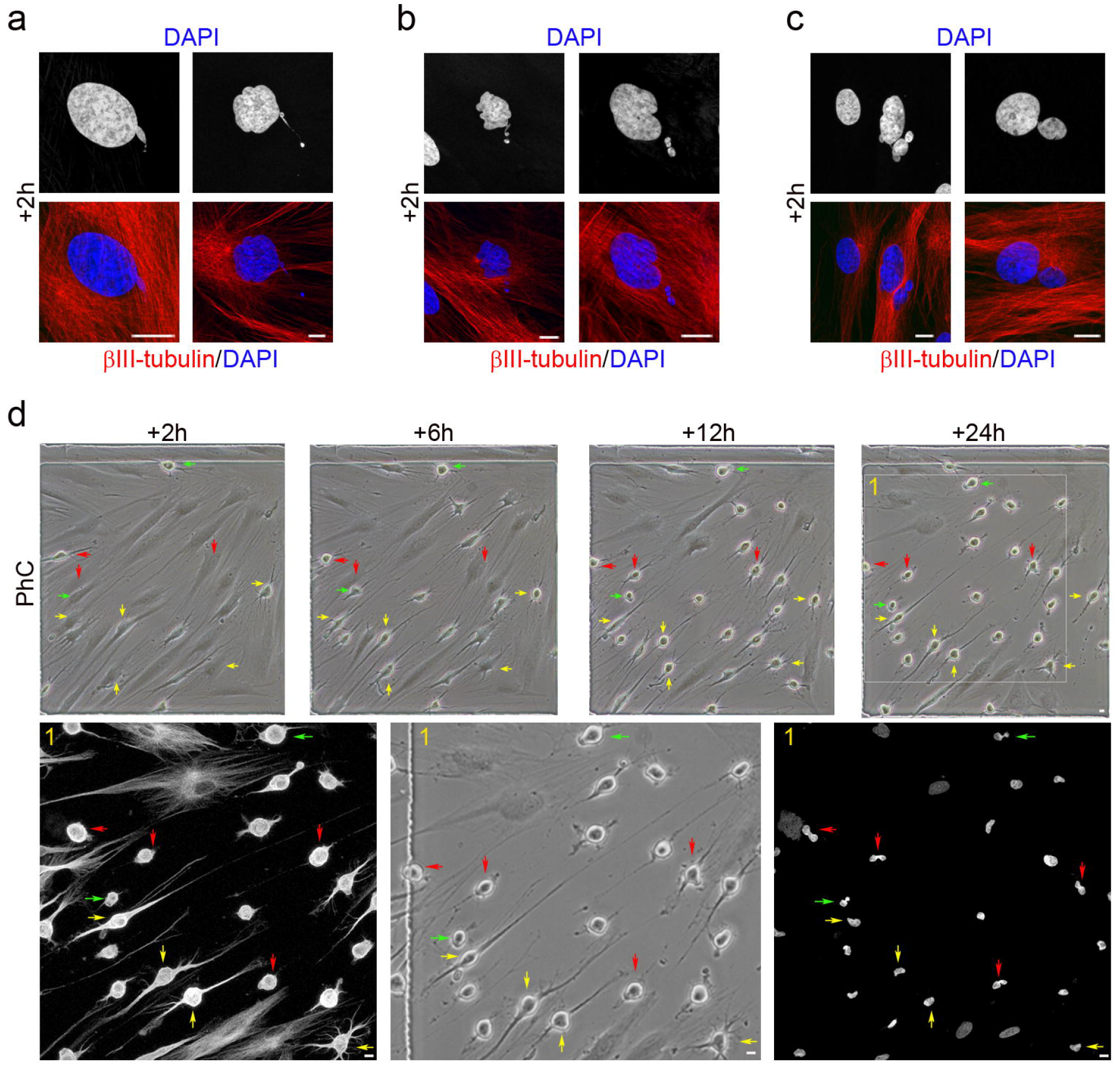
Nuclear shape remodelling occurs during neurogenesis from hBMSCs. DNA-containing structures arose from the main nuclei (**a**) and started to move (**b**) towards specific positions within the cell and temporarily formed lobed nuclei (**c**). These lobed nuclei then connected together through small DNA-containing structures to form nucleoplasmic bridges (**d**, red arrows). Finally, the lobed nuclei connected by internuclear bridges (**d**, green arrows) joined to form single nuclei with an eccentric position within hPDLSC-derived neuron-like cells (**d**, yellow arrows). Scale bar: 10 μm. PhC: Phase-contrast photomicrographs.

**Figure 8.**
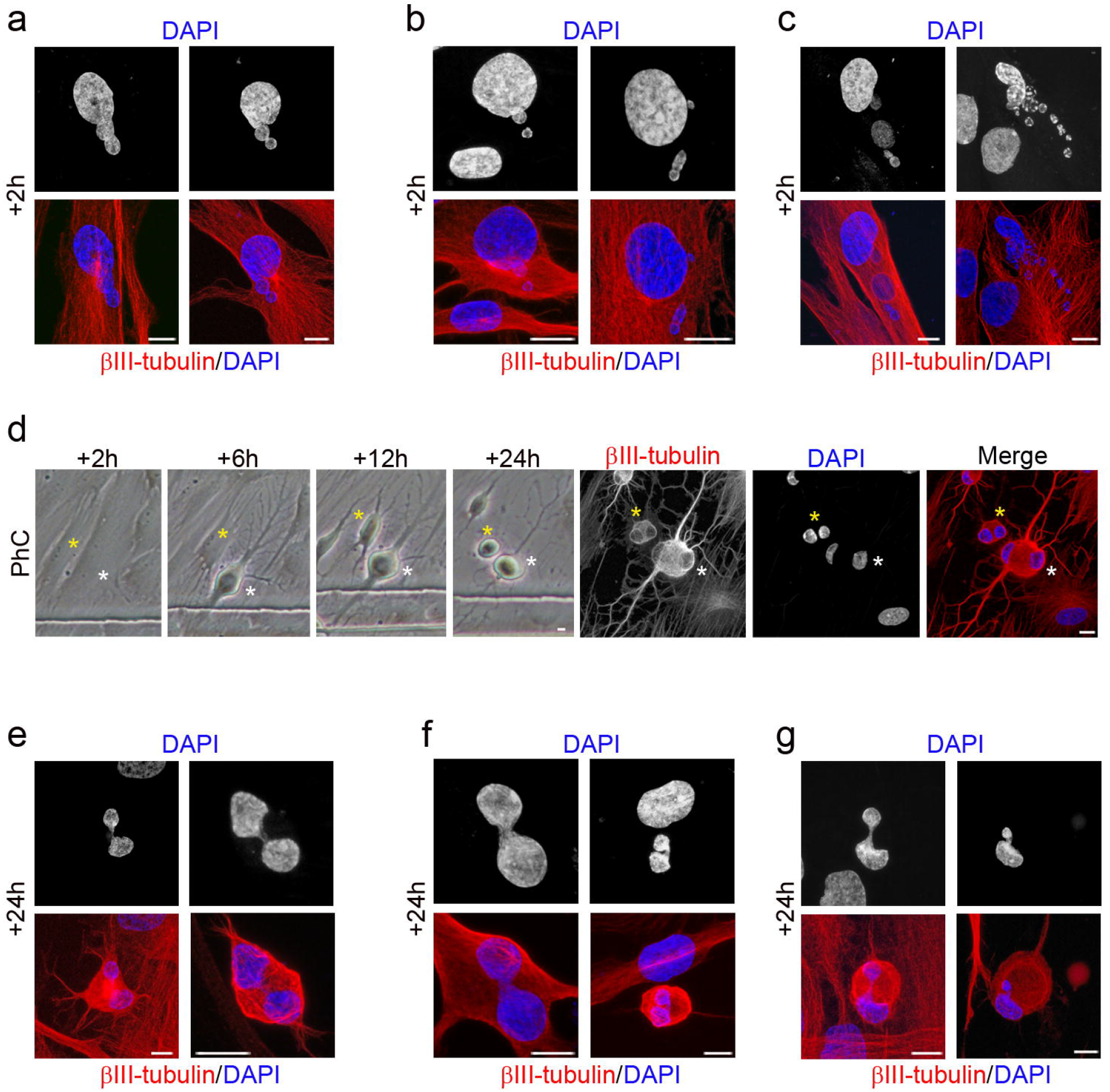
Nuclear shape remodelling occurs during neurogenesis from hDPSCs. DNA-containing structures arose from the main nuclei (**a**) and started to move (**b**) towards specific positions within the cell (**c**) and temporarily formed lobed nuclei (**d**). These lobed nuclei then connected together through small DNA-containing structures to form nucleoplasmic bridges (**e**). Finally, the lobed nuclei connected by internuclear bridges (**f**) joined to form single nuclei with an eccentric position within hPDLSC-derived neuron-like cells (**g**). Scale bar: 10 μm. PhC: Phase-contrast photomicrographs.

Within 24 h of exposure to the neural induction medium we observed lobed nuclei in hDPSCs (Fig. 8d), lobed nuclei connecting together to form nucleoplasmic bridges in hDPSCs (Fig. 8e), lobed nuclei connected by internuclear bridges in hPDLSCs (Fig. 6d, red arrows), hBMSCs (Fig. 7d, red arrows) and hDPSCs (Fig. 8f), and lobed nuclei connected by internuclear bridges merging into a single nucleus in hPDLSCs (Fig. 6d, green arrows), hBMSCs (Fig. 7d, green arrows) and hDPSCs (Fig. 8g). No unusual nuclear structures were observed as hPDLSC-derived (Fig. 6d, yellow arrows) and hBMSC-derived neuron-like cells (Fig. 7d, yellow arrows) gradually acquired a more mature neuron-like morphology.

The results indicate that nuclear remodelling also occurred during *in vitro* neurogenesis from hBMSCs and hDPSCs. hBMSCs and hDPSCs also exhibited unusual nuclear structures and extranuclear bodies in the cellular cytoplasm when they approach a near-spherical shape. These DNA-containing structures displayed a spherical or ovoid shape (Fig. 9a) and some of them appeared to be connected to the main body of the nucleus by thin strands of nuclear material (Fig. 9b, yellow arrows). The hPDLSCs, hBMSCs and hDPSCs had very similar nuclear morphologies during *in vitro* neurogenesis. These results confirm our previous data^24,32^ and further support the idea that hBMSCs, hPDLSCs and hDPSCs follow similar morphological neural development sequences *in vitro*.

**Figure 9.**
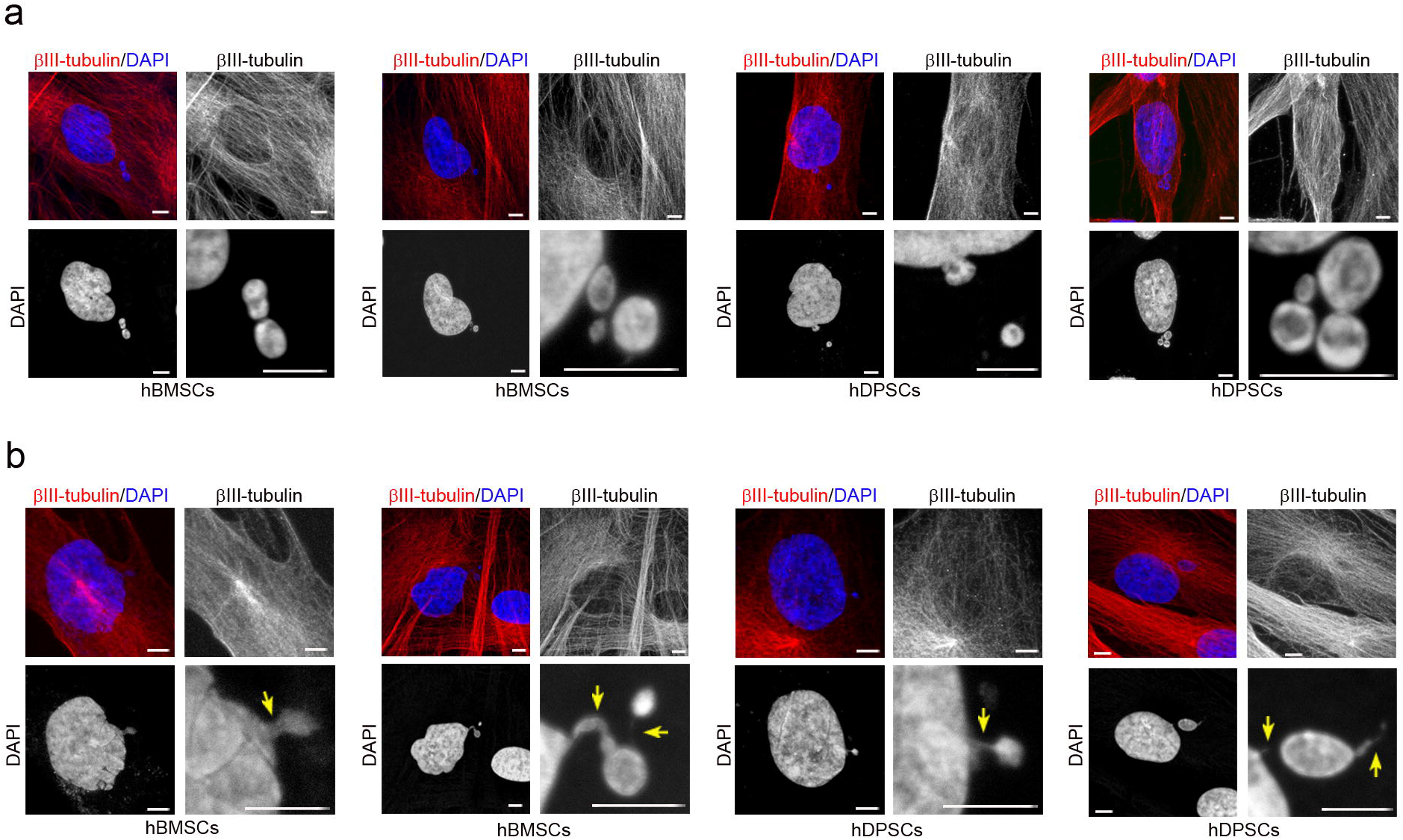
Cytoplasmic DNA-containing structures. Cytoplasmic DNA-containing structures with a spherical or ovoid shape (**a**) and some appeared to be connected to the main body of the nucleus by thin strands of nuclear material (**b**, yellow arrows). Scale bar: 5 μm.

## Discussion

In this study, we have shown that hPDLSCs, hBMSCs and hDPSCs can directly differentiate into neuron-like cells without passing through a mitotic stage. The differentiation of hPDLSCs, hBMSCs and hDPSCs towards a neural lineage occurs through a de-differentiation step (neurogenesis) before the cells re-differentiate to neural phenotypes (neuronal polarisation).

In the present work, we have demonstrated that hPDLSC-derived, hBMSC-derived and hDPSC-derived neuron-like cells produce neurites that gradually adopt a complex morphology, acquiring dendrite-like and axon-like identities. Therefore, our results definitively establish that the rapid acquisition of a neuron-like morphology during neural differentiation occurs though a differentiation trait and is not just an artefact.

Although further research is required to confirm the successful differentiation of hPDLSCs, hBMSCs and hDPSCs into a neural lineage, and finally facilitate the production of autologous cells for cell replacement in a diseased central nervous system, our results provide additional evidence that cells derived from human adult tissues can be converted into neuronal cells without genetic manipulation^33^.

In a previous publication^24^, we reported that nuclear remodelling occurred during *in vitro* neurogenesis from hPDLSCs. We observed that many hPDLSCs had an unusual nuclear structure and chromatin-containing bodies in the cellular cytoplasm when they took on a near-spherical shape. In the present study, we have shown that nuclear remodelling also occurred during *in vitro* neurogenesis from hBMSCs and hDPSCs. We observed that hBMSCs and hDPSCs exhibited unusual nuclear structures and chromatin-containing bodies in the cellular cytoplasm when they became near-spherical in shape. It is important to mention that the hPDLSCs, hBMSCs and hDPSCs had a very similar nuclear morphology during *in vitro* neurogenesis. These results confirm our previous findings^24,32^ and further support that hPDLSCs, hBMSCs and hDPSCs follow similar morphological neural development sequences *in vitro*.

These unusual nuclear structures and extranuclear bodies were described independently in mammals (including in most blood and immune cells^34^), vertebrates, invertebrates, plants and protozoa^34,35,36^. However, the exact role of these extranuclear bodies and their possible correlation with unusual nuclear structures is not yet known^37^. Our results suggest these unusual nuclear structures and extranuclear bodies may be associated with nuclear movement within the cell. Although future analysis involving live-cell nucleus fluorescence labelling and time-lapse microscopy is necessary to determine whether chromatin-containing bodies move within the cell, and if there is any correlation between the formation of these extranuclear bodies and the unusual nuclear structures, a recent study has demonstrated that chromatin-containing bodies arise from the main nuclei, move within the cell and are ultimately loaded in exosomes^38^.

Importantly, the nuclear morphology of hPDLSCs, hBMSCs and hDPSCs observed during the dedifferentiation step (neurogenesis) bears great similarities to the nuclear morphology of neural stem cells located in the ventricular-subventricular zone of the anterolateral ventricle wall in the human foetal brain^39^ and adult mouse brain^40-42^; their nuclear morphology is also very similar to that of many cultured hippocampal neurons^43^. Although it has been suggested that these unusual nuclear structures are associated with quiescence in adult neural stem cells (aNSCs)^42^, our results suggest that they may be associated with nuclear movement within the cell during the initial phases of neurogenesis, but without any relation to cell division.

It has generally been believed that adult neurogenesis occurs progressively through sequential phases of proliferation and neuronal differentiation of aNSCs^44^. However, the current models of aNSC maintenance and differentiation are still contentious because the approaches used to probe stem cell division and differentiation are inherently limited^45^.

It is important to note that very few reports describing newborn neurons in the adult brain have found mitotic chromosomes^46-54^, which would confirm that adult neurogenesis occurs progressively through sequential phases of proliferation. What is more, several studies indicate that some observations interpreted as cell division could be false positives^55-58^. The findings mentioned above and the similarities between the nuclear morphology of hPDLSCs, hBMSCs and hDPSCs during *in vitro* neurogenesis compared to that of aNSCs located in neurogenic niches and cultured hippocampal neurons suggest that aNSCs may also convert into neurons without involving cell division. Future research is necessary to determine the likelihood of this conjecture.

As noted above, there is increasing evidence to suggest that adult cells can assume new fates without asymmetric cell division through dedifferentiation and transdifferentiation processes. These findings call into question the rules of irreversibility and germ-layer restriction as well as the classical definition of a stem cell^4-9,59^. Our results demonstrate that hPDLSCs, hBMSCs and hDPSCs retain plasticity. Therefore, these cells could also help increase our understanding of the mechanisms underlying changes in cell fate and shed light on whether differentiation, de-differentiation and transdifferentiation occur in different contexts or are different terms for the same cellular process.

## Methods

### Ethical conduct of research

The authors declare that all protocols used to obtain and process all human samples were approved by the local ethics committees (UMH.IN.SM.03.16, HULP3617.05/07/2012 and HUSA19/1531.17/02/2020) according to Spanish and European legislation and conformed to the ethical guidelines of the Helsinki Declaration. Donors provided written informed consent before obtaining samples.

### Isolation and culture of hPDLSCs and hDPSCs

To obtain hPDLSCs, human premolars were extracted and collected from three different healthy adult donors undergoing orthodontic therapy in Murcia dental hospital (Spain). Human periodontal ligament (hPDL) was scraped from the middle third region of the root surface. To obtain hPDSCs, human third molars were extracted and collected from three different healthy adult donors undergoing orthodontic therapy in Murcia dental hospital (Spain). Human dental pulp (hDP) was harvested from the teeth.

Dissociated cell cultures of hPDL and hDP tissues were prepared as previously described^24,60^. After washing the extracted hPDL and hDP with Ca and Mg-free Hank’s balance salt solution (HBSS; Gibco), hPDL and hDP was digested with 3 mg/ml type I collagenase (Worthington Biochemical Corporation) and 4 mg/ml dispase II (Gibco) in alpha modification minimum essential medium eagle (α-MEM; Sigma-Aldrich) for 1 h at 37 °C. The reaction was stopped by the addition of α-MEM. hPDL and hDP derived from different subjects were pooled together to obtain single cell suspensions by passing the cells through a 70 μm strainer (BD Falcon). Cells were centrifuged, and the pellet was resuspended in serum-containing media (designated as the basal media), composed of DMEM low glucose medium (Thermo Fisher Scientific) supplemented with 10% fetal bovine serum (Sigma-Aldrich), non-essential amino acid solution (Sigma-Aldrich), 100 units/ml penicillin-streptomycin (Sigma-Aldrich) and 2 mM l-glutamine (Sigma-Aldrich). The cell suspension was plated into six-well multiwell plates (BD Falcon) and incubated at 37 °C in 5% CO2. All studies were performed using hPDLSCs and hDPSCs expanded within culture passages 3-4.

### Isolation and culture of hBMSCs

Bone marrow aspirates were obtained by percutaneous direct aspiration from the iliac crest of 5 healthy volunteers at University Hospital Virgen de la Arrixaca (Murcia, Spain). Bone marrow was collected with 20 U/ml sodium heparin, followed by a Ficoll density gradient-based separation by centrifugation at 540g for 20 min. After, mononuclear cell fraction was collected, washed twice with Ca^2+^/Mg^2+^-free phosphate buffered saline (PBS) (Gibco Invitrogen) and seeded into 175-cm2 culture flasks (Nunc, Thermo Fisher Scientific) at a cell density 1.5×10^5^ cells/cm^2^ in DMEM low glucose medium (Thermo Fisher Scientific) supplemented with 10% fetal bovine serum (FBS; Lonza), 1% GlutaMAX (Thermo Fisher Scientific) and 1% penicillin/streptomycin (Thermo Fisher Scientific). After 3 days of culture at 37°C and 5% CO_2_, non-attached cells were removed and fresh complete medium was added. Culture media were renewed every 2 days, and the isolated hBMSCs were passaged when cultures were 70-80% confluent. All studies were performed using hBMSCs expanded within culture passages 3-4.

### Time-lapse microscopy of hPDLSCs, hPDSCs and hBMSCs cultured in neural induction media

We used μ-Dish 35 mm, high Grid-500 (Ibidi) for live cell imaging. Numeric marks on the bottom of each dish allow users to identify the location of cells. Cells were plated onto collagen IV (Sigma-Aldrich) coated plastic or glass coverslips, and maintained in basal media or neural induction media.

To induce neural differentiation, cells at passage 3–4 were allowed to adhere to the plates overnight. Media was removed the following day and the cells were cultured for 2 days in serum-free media (designated as the neural basal media) consisting in Dulbecco’s modified Eagle’s medium/F12 (DMEM/F12 Glutamax, Gibco) supplemented with N2-supplement (R&D systems), 0.6% glucose (Sigma-Aldrich), 5mM HEPES (Sigma-Aldrich), 0.5% human serum albumin (Sigma-Aldrich), 0.0002% heparin (Sigma-Aldrich), non-essential amino acid solution (Sigma-Aldrich) and 100 units/ml penicillin-streptomycin (Sigma-Aldrich). On day 3, cells were cultured in neural induction media, consisting in the neural basal media supplemented with 500nM retinoic acid (Sigma-Aldrich), 1mM dibutyryl cAMP (Sigma-Aldrich) and growth factors like BDNF (10□ng/ml; Peprotech), GDNF (10□ng/ml; Peprotech) and IGF-1 (10□ng/ml; R&D systems). We perform time-lapse phase-contrast microscopy within the first 24 hr after neural induction media was added directly to the cells.

### Immunocytochemistry

A standard immunocytochemical protocol was used as previously described^24,60^. Cells were plated onto collagen IV (Sigma-Aldrich) coated plastic or glass coverslips, and maintained in basal media or neural induction media. Cells were rinsed with PBS and fixed in freshly prepared 4% paraformaldehyde (PFA; Sigma-Aldrich). Fixed cells were blocked for 2 h in PBS containing 10% normal horse serum (Gibco) and 0.25% Triton X-100 (Sigma) and incubated overnight at 4 °C with antibodies against β-III-tubulin (TUJ1; 1:500, Covance) in PBS containing 1% normal horse serum and 0.25% Triton X-100. On the next day, cells were rinsed and incubated with the secondary antibody: Alexa Fluor® 594 (anti-mouse; 1:500, Molecular Probes). Cell nuclei were counterstained with DAPI (0.2 mg/ml in PBS, Molecular Probes). Alexa Fluor 488® phalloidin was used to selectively stains F-actin (Molecular Probes). Data are representative of ten independent experiments per condition.

### Images and Data Analyses

Analyses and photography of visible and fluorescent stained samples were carried out in an inverted Leica DM IRB microscope equipped with a digital camera Leica DFC350FX (Nussloch) or in confocal laser scanning microscope Leica TCS-SP8. Digitized images were analyzed using LASX Leica confocal software. We used Photoshop software to improve the visibility of fluorescence images without altering the underlying data.

## Author Contributions

C.B. conceived of the study, designed the study, carried out the molecular lab work and drafted the manuscript. M.M. and D.G. carried out the molecular lab work and participated in data analysis. J.M. helped draft the manuscript and financial support. S.M. conceived of the study, helped draft the manuscript and financial support. All authors discussed and commented on the manuscript.

## Competing Interest

The other authors declare no competing interests.

## Funding

This work has been funded by Instituto de Salud Carlos III through the project “RD16/001/0010” (Co-funded by European Regional Development Fund/European Social Fund “Investing in your future”) and Spanish MINECO/AEI/FEDER (SAF2017-83702-R).

## Figure legends

**Figure S1.**
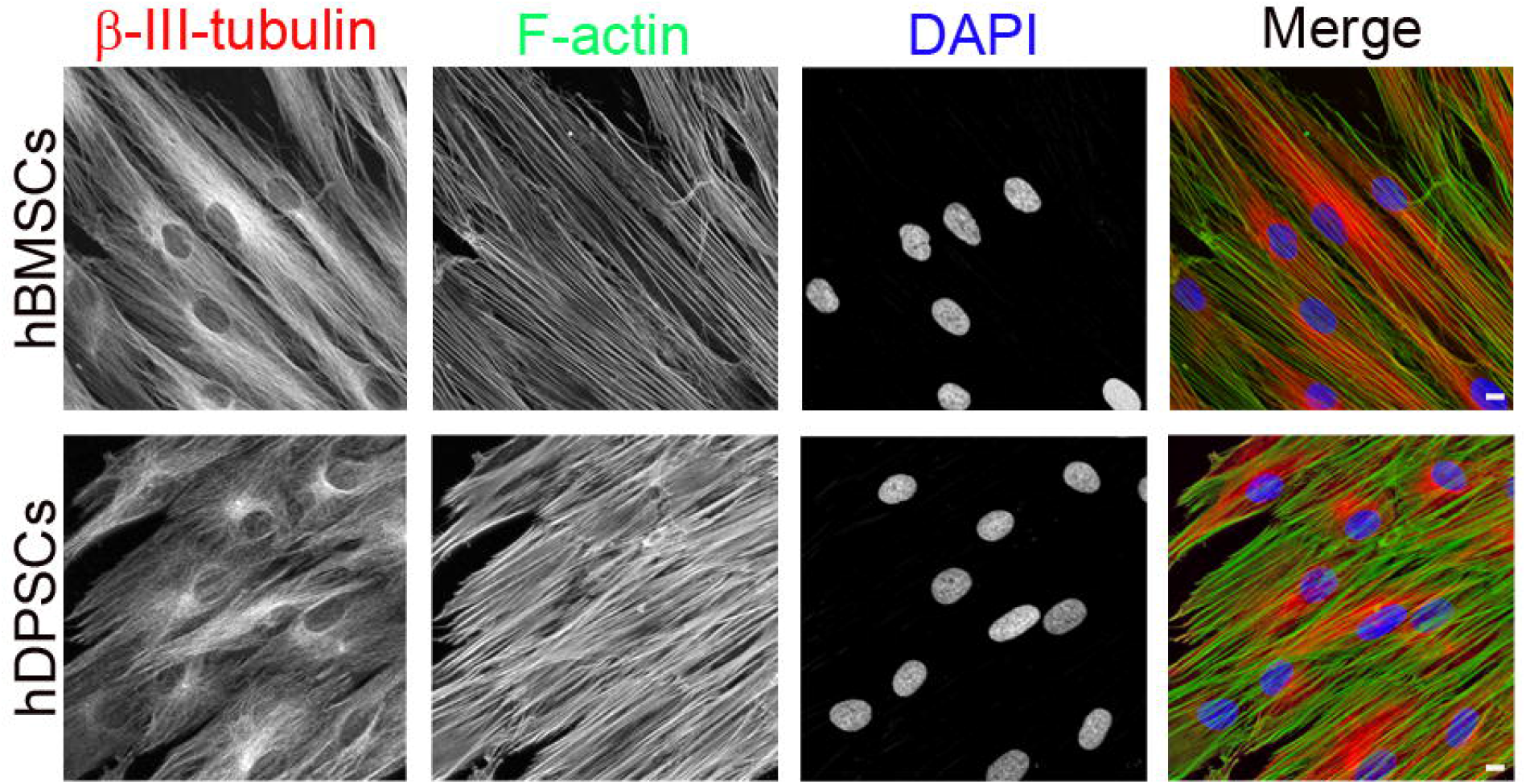
Morphology of hBMSCs and hDPSCs cultured in basal media. During interphase, undifferentiated hBMSCs and hDPSCs presented a fibroblast-like morphology with actin microfilaments and β-III tubulin microtubules oriented parallel to the cell’s longitudinal axis and an ellipsoidal nucleus, often located in the centre of the cell. Scale bar: 10 μm.

## Notes

### Competing Interest Statement

The authors have declared no competing interest.

